## Supplementary Methods for "Weight Pulling: A Novel Mouse Model of Human Progressive Resistance Exercise": Supplemental Methods.pdf

CellProfiler version 4.1.3 was downloaded from https://cellprofiler.org/previous-releases and used on a 64-bit Windows 10 PC (Intel Core i7-9850H CPU @ 2.60 GHz, 64 GB RAM). The fiber cross-sectional area CellProfiler pipeline and myonuclei CellProfiler pipeline, as well as sample images for running these pipelines, are provided within the supplementary methods zip file. The CellProfiler pipelines represent modifications of the original Muscle2View pipeline [1]. Before attempting to employ these pipelines the reader should be familiar with the manuscript which describes the Muscle2View pipeline and the CellProfiler manual (module section, https://cellprofiler-manual.s3.amazonaws.com/CellProfiler-4.1.3/index.html).

Fiber Cross-Sectional Area Pipeline

This pipeline was used to obtain measurements of the whole muscle mid-belly cross-sectional area (CSA), mean CSA for all fibers, the total number of fibers per cross-section, fiber type-specific CSA, the aspect ratio of the fibers (i.e., max ferret diameter / min ferret diameter) and the number of fibers per cross-section that belonged to each fiber type. The pipeline consists of 23 sequential modules which can be run without the need for human supervision and generally takes less than 2 minutes per sample to run to completion.

*Image Load*

Initially, mid-belly cross-section images corresponding to Laminin, Type I, Type IIA, and Type IIB MHC were loaded as 16-bit TIFF files into the file list area under the Images module. Each image was extracted from the file name using the Metadata module and the regular expression *Ch(?P<ChannelNumber>\d+)*. Next, images were grouped using the NamesAndTypes module based on their <ChannelNumber>. For instance, we named Ch0 as "Laminin", Ch1 as "Type IIB", Ch2 as "Type IIA", and Ch3 as "Type I". Samples images with the names Ch0, Ch1, Ch2 and Ch3 are provided in the "Fiber Cross-Sectional Area Pipeline" folder that is contained within the supplemental methods zip file. These images can be directly loaded into the CellProfiler user interface and used to test the functionality of the pipeline.

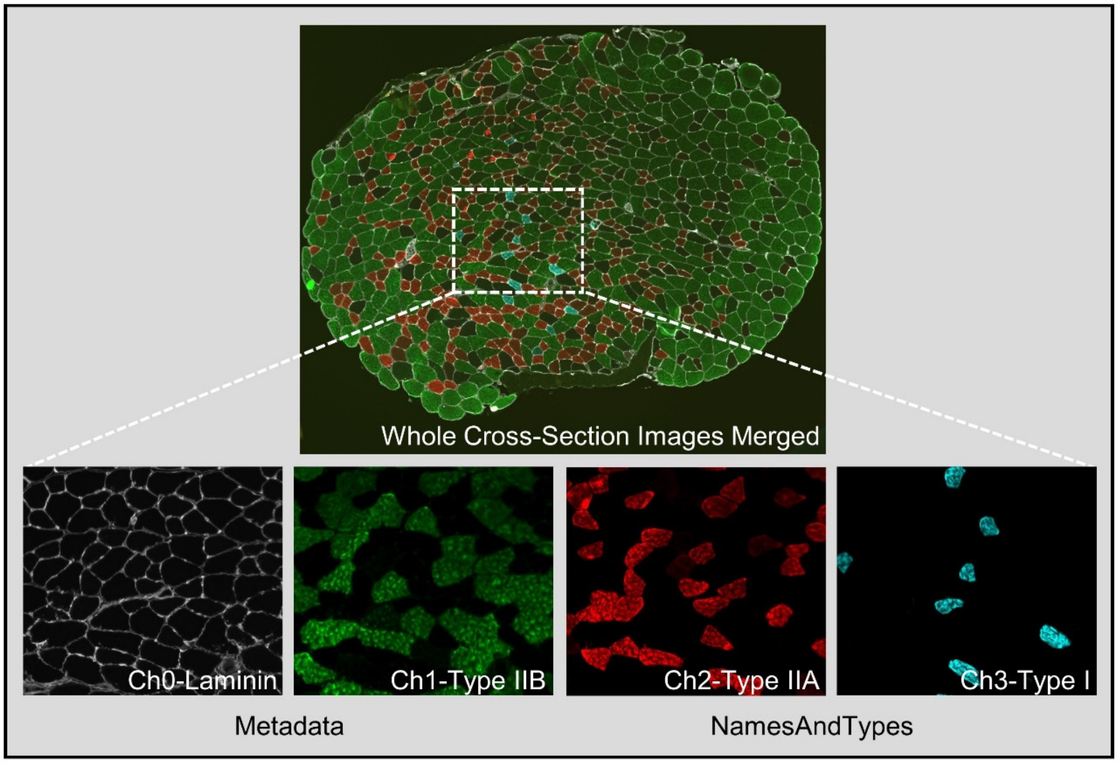

*Laminin Image Processing*

To accurately identify individual fibers, the Laminin image was optimized through six modules (MedianFilter, Closing, EnhanceOrSupressFeatures, ImageMath, Opening, ImageMath) as described in Muscle2View [1].

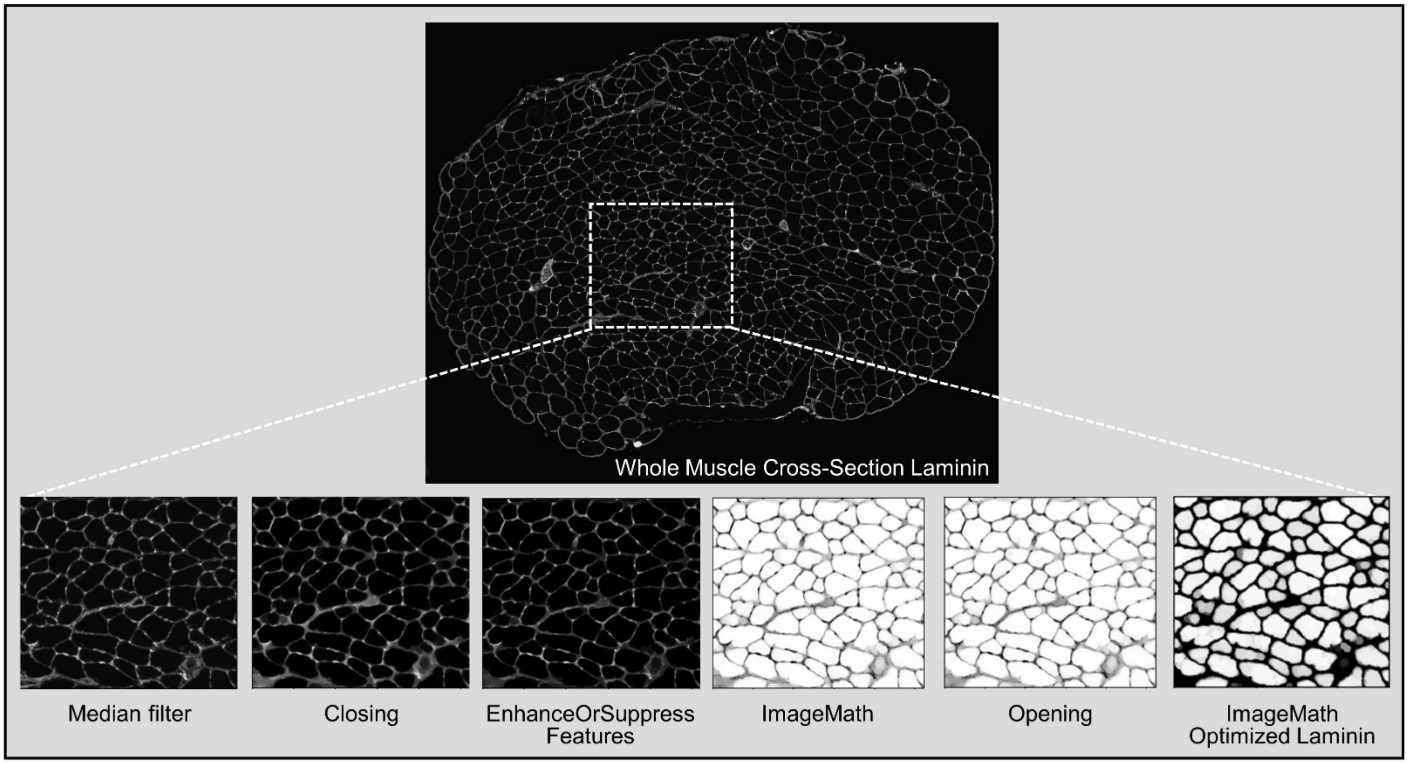


*Muscle Fiber Segmentation and Fiber Type Characterization*

The IdentifiedPrimaryObjects module was used to identify muscle fibers by using the optimized Laminin image created from ImageMath module. The FilterObjects module with settings of 0.93 Eccentricity, and 0.5 FormFactor were used to filter out extremely oblique / longitudinally oriented fibers and inappropriately shaped fibers, respectively. MedianFilter and Threshold modules were used to get optimized fiber type-specific images prior to applying the MaskObjects module. The threshold module produces a binary (i.e., black and white) image based on a threshold that can be pre-selected or calculated automatically using one of many available methods (e.g., Minimum Cross-Entropy, Otsu, Robust Background). After the threshold value has been determined, the Threshold module will set pixel intensities below the value to zero (black) and above the value to one (white). Therefore, it is important to pay attention to the selection of the threshold method to avoid high background pixel intensities. Specific fiber types were distinguished by using the MaskObjects module. This module allows you to delete the objects or portions of objects that are outside of a region (mask) that you specify. If using a masking image, the mask is composed of the foreground (white portions); if using a masking object, the mask is composed of the area within the object. You can choose to remove only the portion of each object that is outside of the region, remove the whole object if it is partially or fully outside of the region, or retain the whole object unless it is fully outside of the region. For instance, the filtered fibers image shown below was used as a masking image (foreground), and the optimized Type I image was used as the masking object. Type I fibers were counted only if the optimized Type I image (masking object) overlapped at least 50% with filtered fibers image (masking image). After counting Type I fibers, the masking image was inverted, which means that the non-type I fibers image was used as the masking object for the Type IIA fibers image. For the Type IIB and IIX fiber identification, the non-Type I + non-Type IIA fiber image was used as the masking image. The inverted optimized Type IIB image was then used as the masking object to identify Type IIX fibers (i.e., fibers that were non-TypeI, non-Type IIA, and non-Type IIB). Finally, the MeasureObjectSizeShape and MeasureImageAreaOccupied modules were used to measure fiber type-specific CSA and mid-belly CSA.

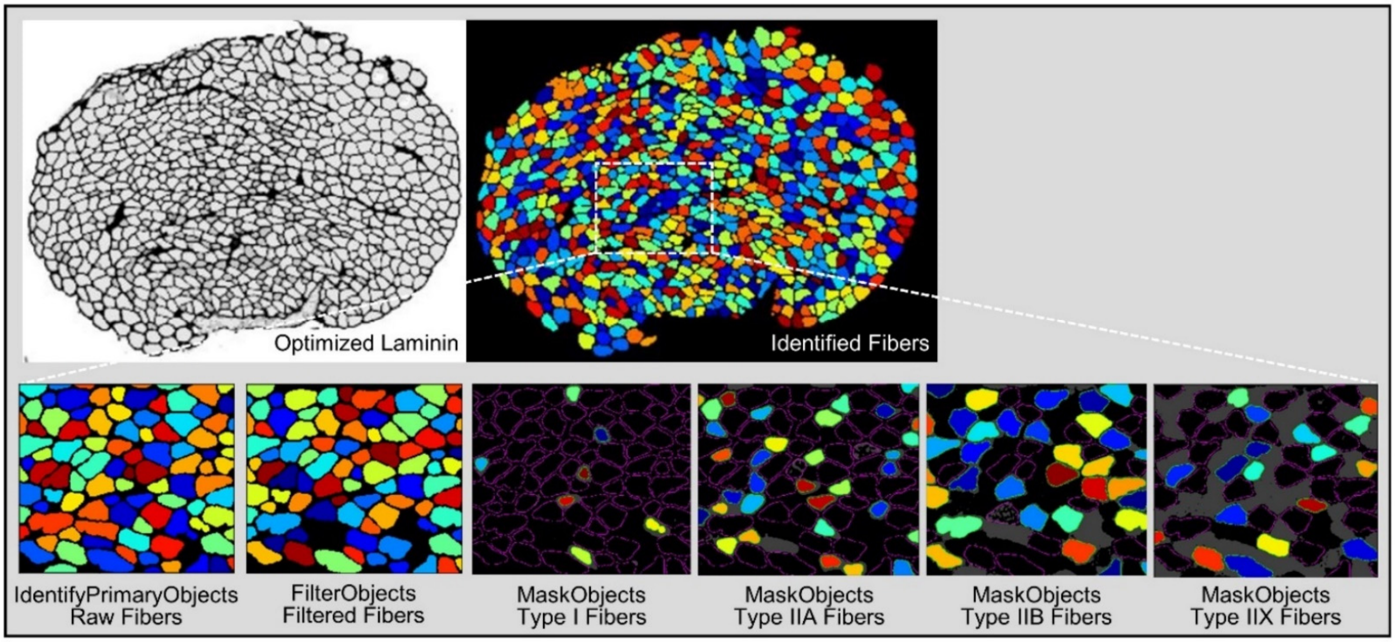


*Export Results*

The ExportToSpreadsheet module allows for the export of the data as combined average values for the entire image, as well as the export of individual values for each type of identified object (e.g., every Type I, IIA, IIX, IIB fiber). The names of these files will be titled as Export_AverageCSA.csv and Export_IndividualFiberCSA.csv, respectively. To export the files correctly, please choose the appropriate output file location that you desire. Also, the export data prefix names in these files are dependent on the selected measurements within the CellProfiler pipeline and the nomenclature may not be intuitively obvious. Hence, to address this, we have provided a table that describes what each of the output values mean. Finally, it is also very important to recognize that all area based measurements are presented in terms of pixel area. For final data analysis, the user will need to convert the pixel area values to real area values.

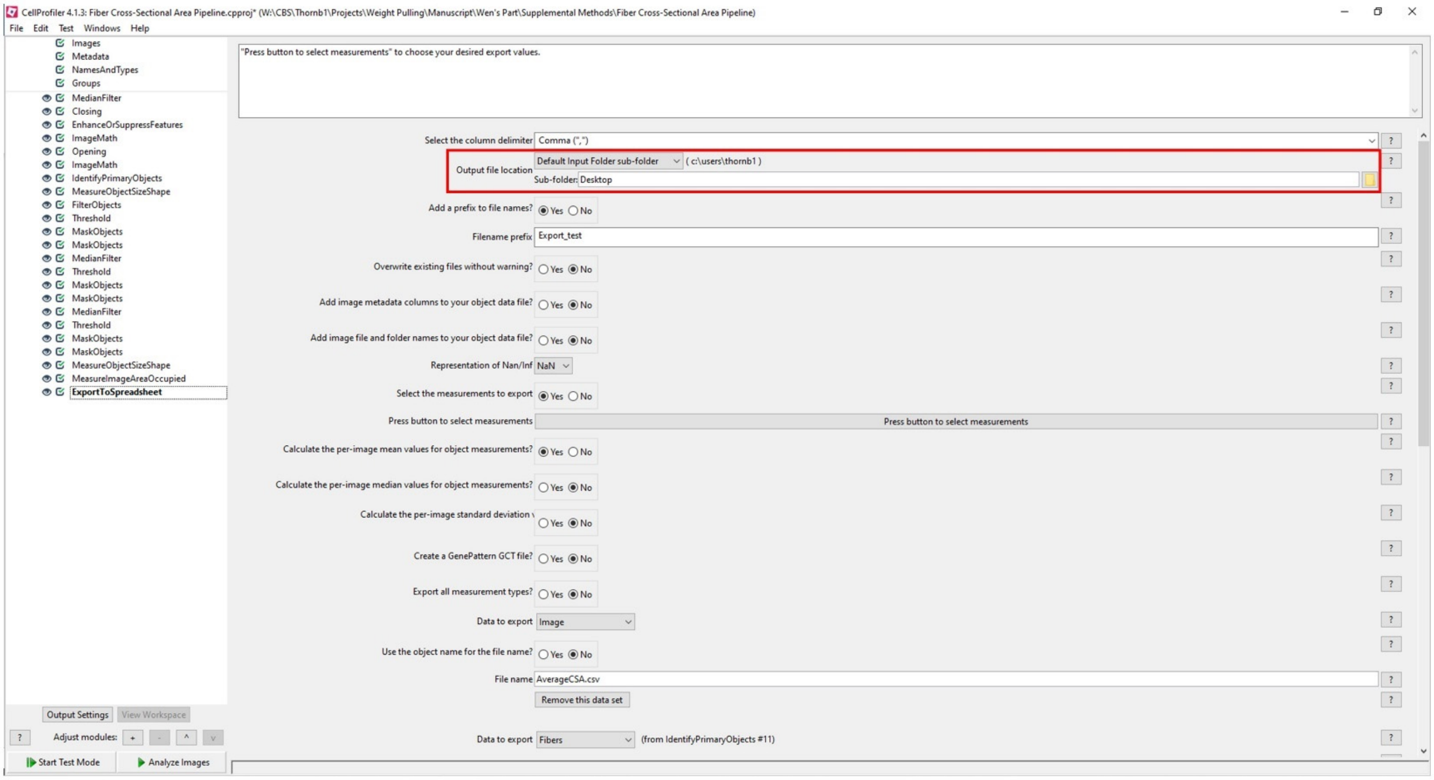

### *Description of the Exported Results*

"FilteredFibers" = All fibers minus those that were filtered out because of the 0.93 Eccentricity and 0.5 FormFactor settings
(i.e., the longitudinally oriented and/or inappropriately shaped fibers)

| | |
|---|---|
| AreaOccupied_AreaOccupied_Fibers | Whole muscle cross-sectional area based on all of the identified fibers |
| AreaOccupied_AreaOccupied_FilteredFibers | Whole muscle cross-sectional area based on all of the fibers from the "FilteredFibers" list |
| Count_Fibers | The total number of fibers based on all of the identified fibers |
| Count_FilteredFibers | The total number of "FilteredFibers" |
| Count_I | The number of Type I fibers from the "FilteredFibers" list |
| Count_IIA | The number of Type IIA fibers from the "FilteredFibers" list |
| Count_IIX | The number of Type IIX fibers from the "FilteredFibers" list |
| Count_IIB | The number of Type IIB fibers from the "FilteredFibers" list |
| Mean_Fibers_AreaShape_Area | Mean fiber cross-sectional area based on all of the identified fibers |
| Mean_FilteredFibers_AreaShape_Area | Mean fiber cross-sectional area based on all of the fibers from the ""FilteredFibers" list |
| Mean_Fibers_AreaShape_MaxFeretDiameter | Mean Maximal Feret Diameter based on all of the fibers |
| Mean_FilteredFibers_AreaShape_MaxFeretDiameter | Mean Maximal Feret Diameter based on all of the fibers from the ""FilteredFibers" list |
| Mean_Fibers_AreaShape_MinFeretDiameter | Mean Minimal Feret Diameter based on all of the fibers |
| Mean_FilteredFibers_AreaShape_MinFeretDiameter | Mean Minimal Feret Diameter based on all of the fibers from the ""FilteredFibers" list |
| Mean_I_AreaShape_Area | Mean Type I fiber cross-sectional area based on all of the Type I fibers from the ""FilteredFibers" list |
| Mean_I_AreaShape_MaxFeretDiameter | Mean Type I Maximal Feret Diameter based on all of the Type I fibers from the ""FilteredFibers" list |
| Mean_I_AreaShape_MinFeretDiameter | Mean Type I Minimal Feret Diameter based on all of the Type I fibers from the ""FilteredFibers" list |
| Mean_IIA_AreaShape_Area | Mean Type IIA fiber cross-sectional area based on all of the Type IIA fibers from the ""FilteredFibers" list |
| Mean_IIA_AreaShape_MaxFeretDiameter | Mean Type IIA Maximal Feret Diameter based on all of the Type IIA fibers from the ""FilteredFibers" list |
| Mean_IIA_AreaShape_MinFeretDiameter | Mean Type IIA Minimal Feret Diameter based on all of the Type IIA fibers from the ""FilteredFibers" list |
| Mean_IIX_AreaShape_Area | Mean Type IIX fiber cross-sectional area based on all of the Type IIX fibers from the ""FilteredFibers" list |
| Mean_IIX_AreaShape_MaxFeretDiameter | Mean Type IIX Maximal Feret Diameter based on all of the Type IIX fibers from the ""FilteredFibers" list |
| Mean_IIX_AreaShape_MinFeretDiameter | Mean Type IIX Minimal Feret Diameter based on all of the Type IIX fibers from the ""FilteredFibers" list |
| Mean_IIB_AreaShape_Area | Mean Type IIB fiber cross-sectional area based on all of the Type IIB fibers from the ""FilteredFibers" list |
| Mean_IIB_AreaShape_MaxFeretDiameter | Mean Type IIB Maximal Feret Diameter based on all of the Type IIB fibers from the ""FilteredFibers" list |
| Mean_IIB_AreaShape_MinFeretDiameter | Mean Type IIB Minimal Feret Diameter based on all of the Type IIB fibers from the ""FilteredFibers" list |
| AreaShape_Area | Cross-sectional area of the specific object (i.e., a single fiber) |
| AreaShape_MaxFeretDiameter | Maximal Feret Diameter of the specific object (i.e., a single fiber) |
| AreaShape_MinFeretDiameter | Minimal Feret Diameter of the specific object (i.e., a single fiber) |

*Myonuclei Pipeline Specifc Output*

| | |
|---|---|
| Count_Nuclei | The total number of nuclei |
| Count_Myonuclei | The total number of myonuclei |
| | *Note: Interstital nuclei are determined by subtracting the number of myonuclei from the total nuclei* |

Myonuclei Pipeline

The myonuclei pipeline is very similar to the fiber cross-sectional area pipeline with respect to the loading of images and processing of the dystrophin image for the subsequent identification of fibers. The only difference is that the myonuclei pipeline does not focus on specific fiber types. Samples images with the names Ch0, and Ch1 are provided in the "Myonuclei Pipeline" folder that is contained within the Supplemental methods zip file. These images can be loaded into the CellProfiler interface and used to test the functionality of the myonuclei pipeline.

*Myonuclei Identification*

The IdentifyPrimaryObjects module is used to identify the region occupied by nuclei and uses a pixel size range of 3-50. The MaskObjects module is used to identify myonuclei based on the overlapping region between the filtered fibers image (masking image) and nuclei (mask object). The FilterObjects module is used to remove the interstitial nuclei and the remaining nuclei are counted as myonuclei.

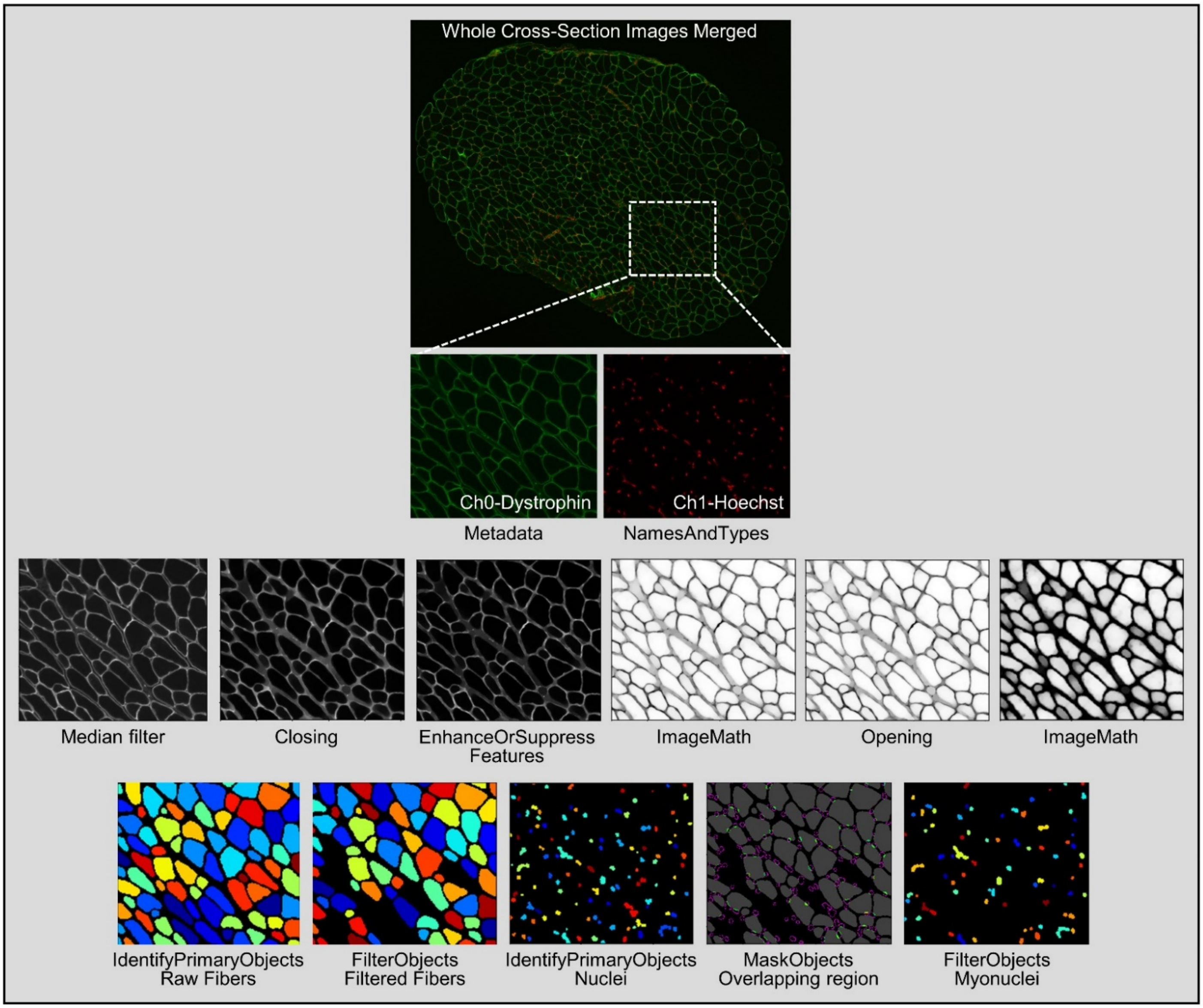


*Export Results*

The ExportToSpreadsheet module allows for the export of the data as a file titled ExportMyonuclei_.csv. To export the file correctly, choose the appropriate output file location as described for the fiber cross-sectional area pipeline and please refer to the above "*Description of the Exported Results*" table for a breif description of what each of the output values mean. Also, bear in mind that all area based measurements are presented in terms of pixel area. For final data analysis, the user will need to convert the pixel area values to real area values.
