## Supplementary figures and images for "Weight Pulling: A Novel Mouse Model of Human Progressive Resistance Exercise"

### Ch0.tif

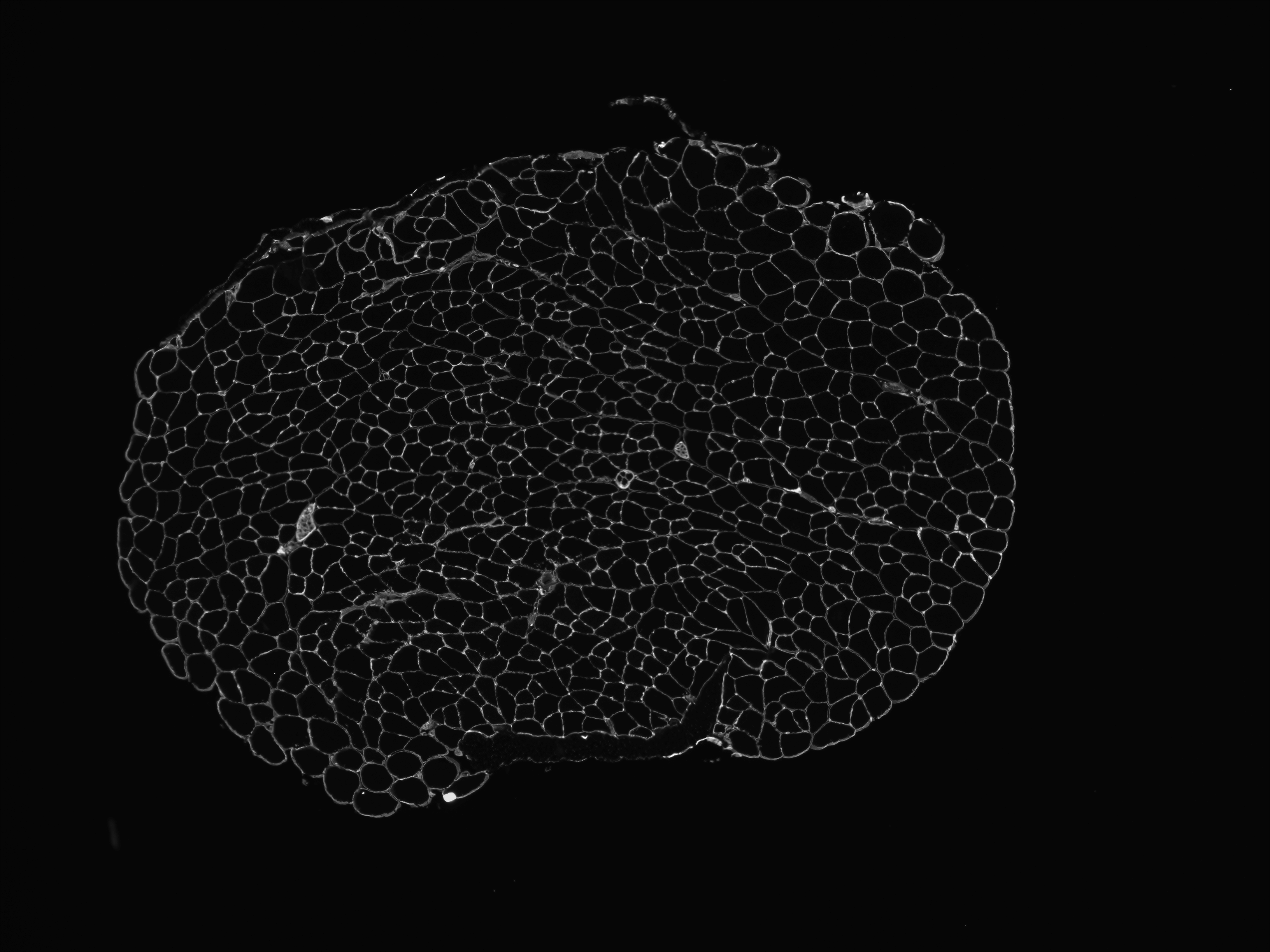

### Ch0.tif

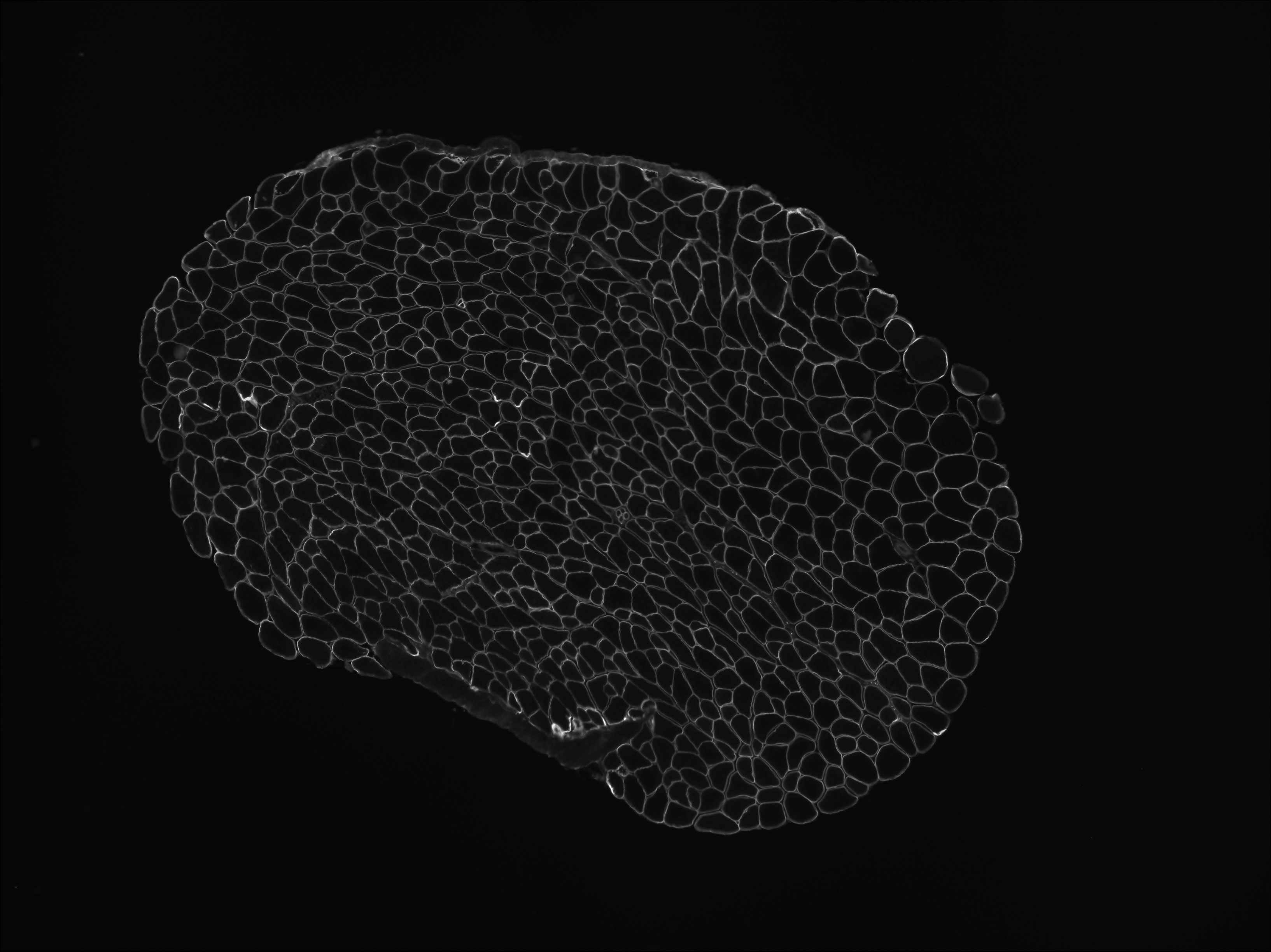

### Ch1.tif

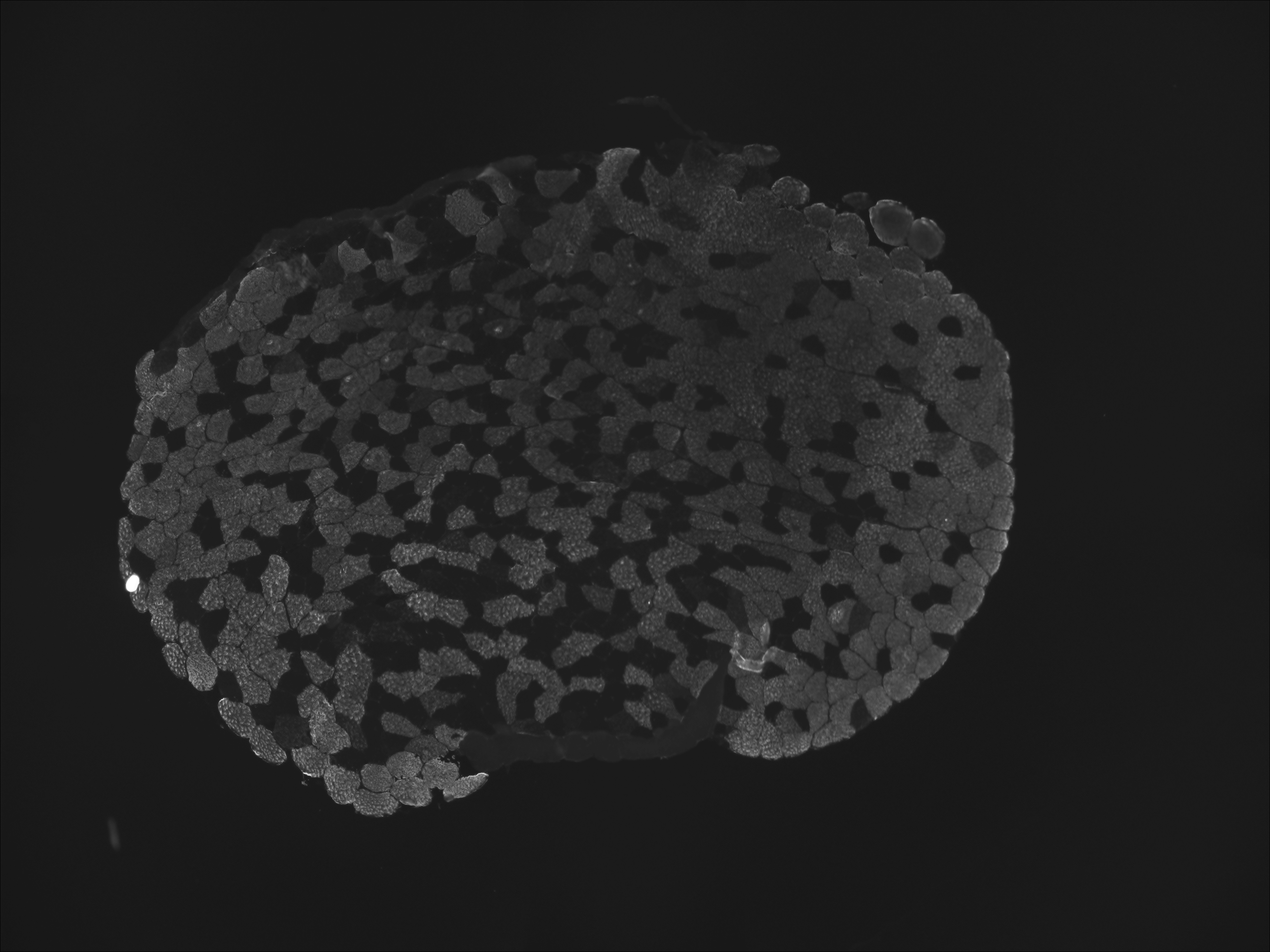

### Ch1.tif

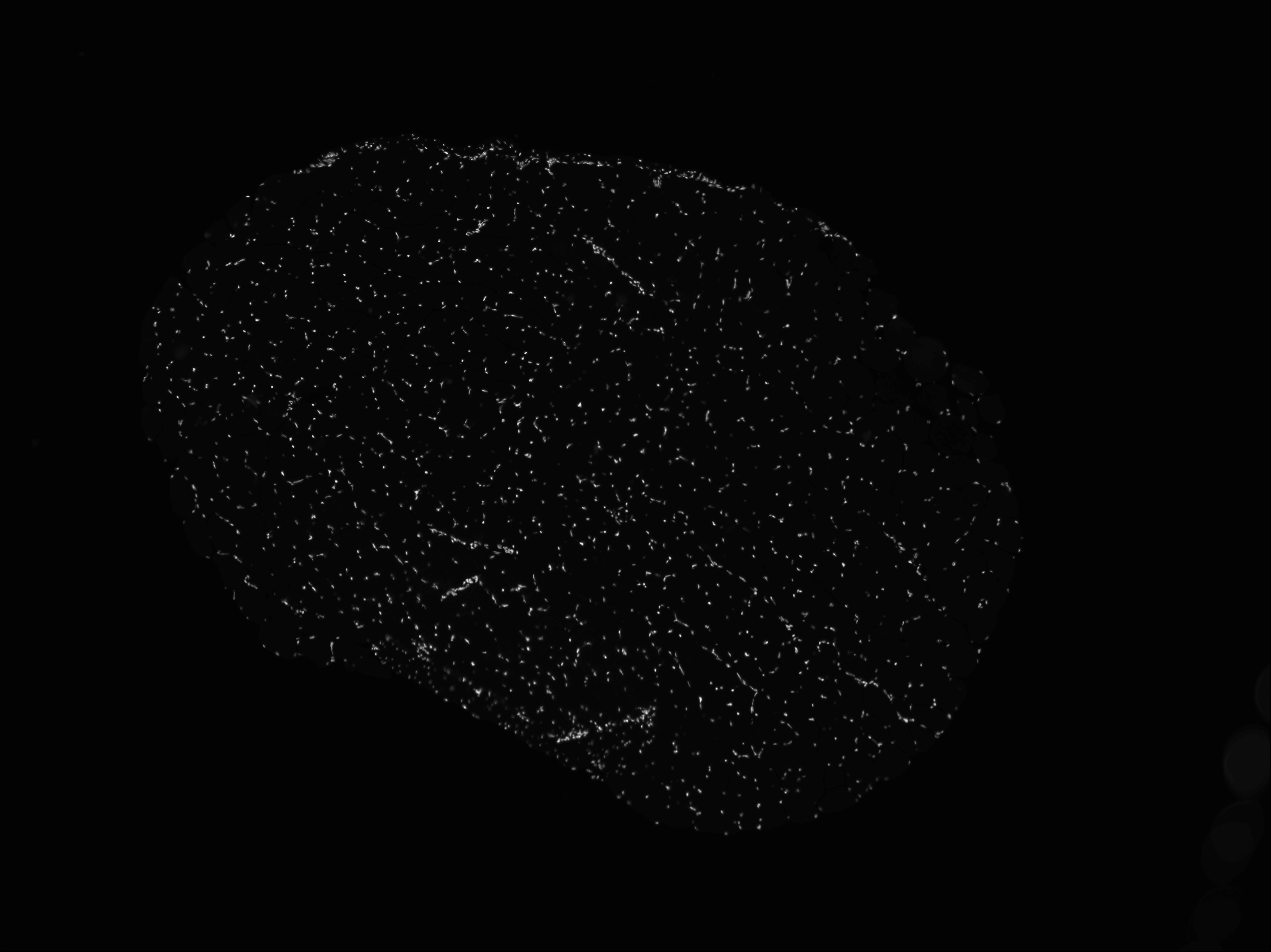

### Ch2.tif

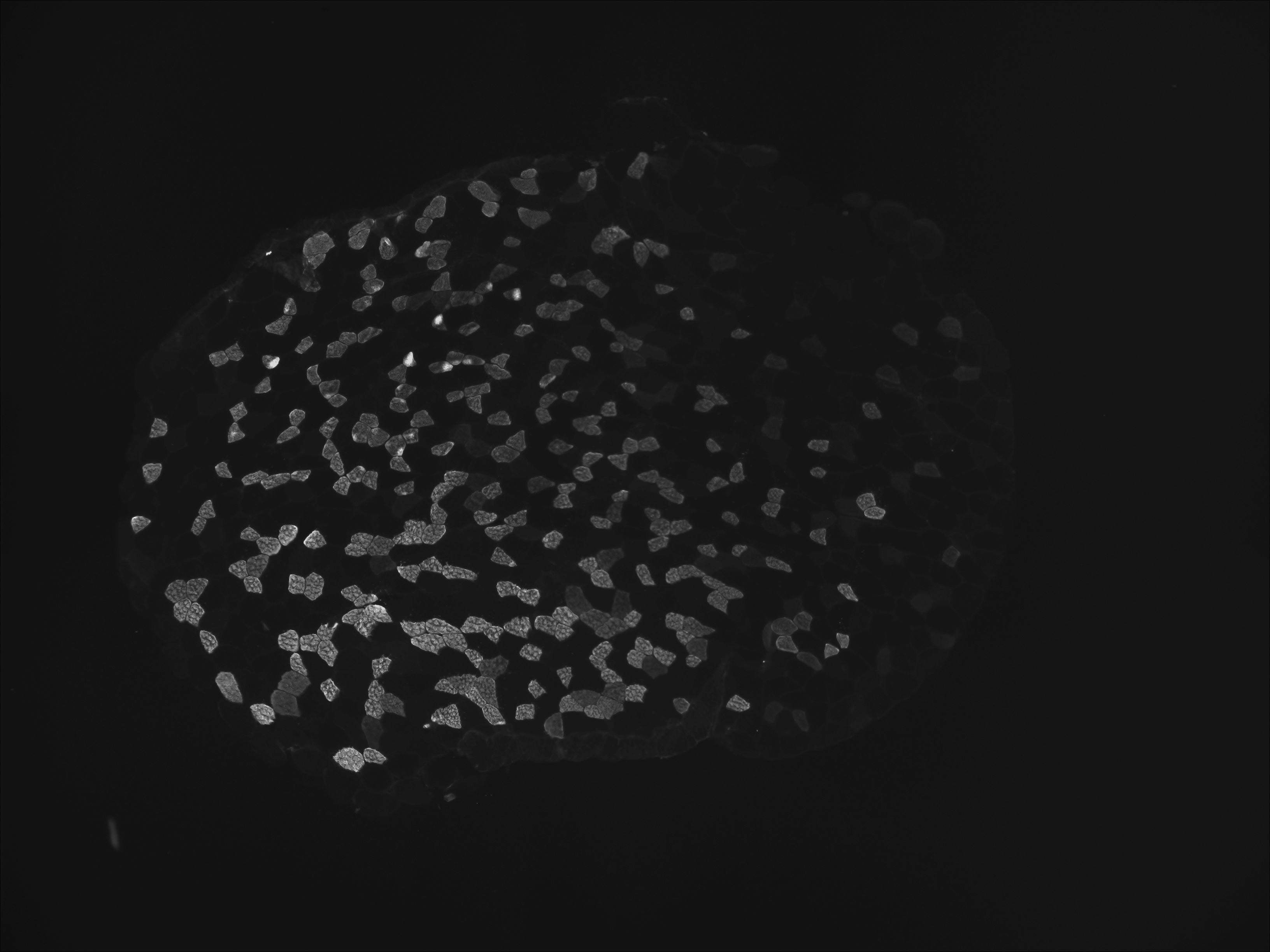

### Ch3.tif

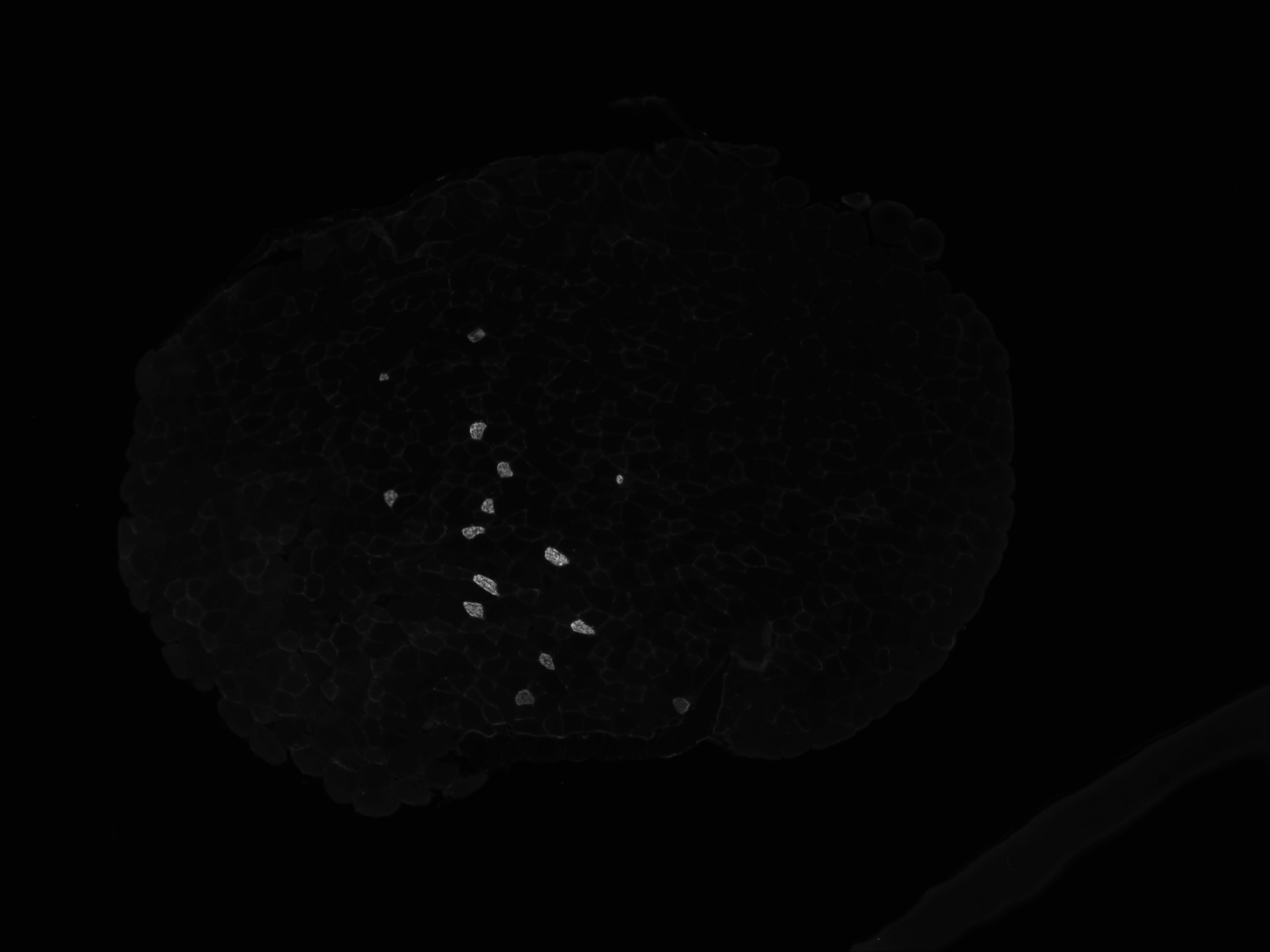

### Graphical Abstract

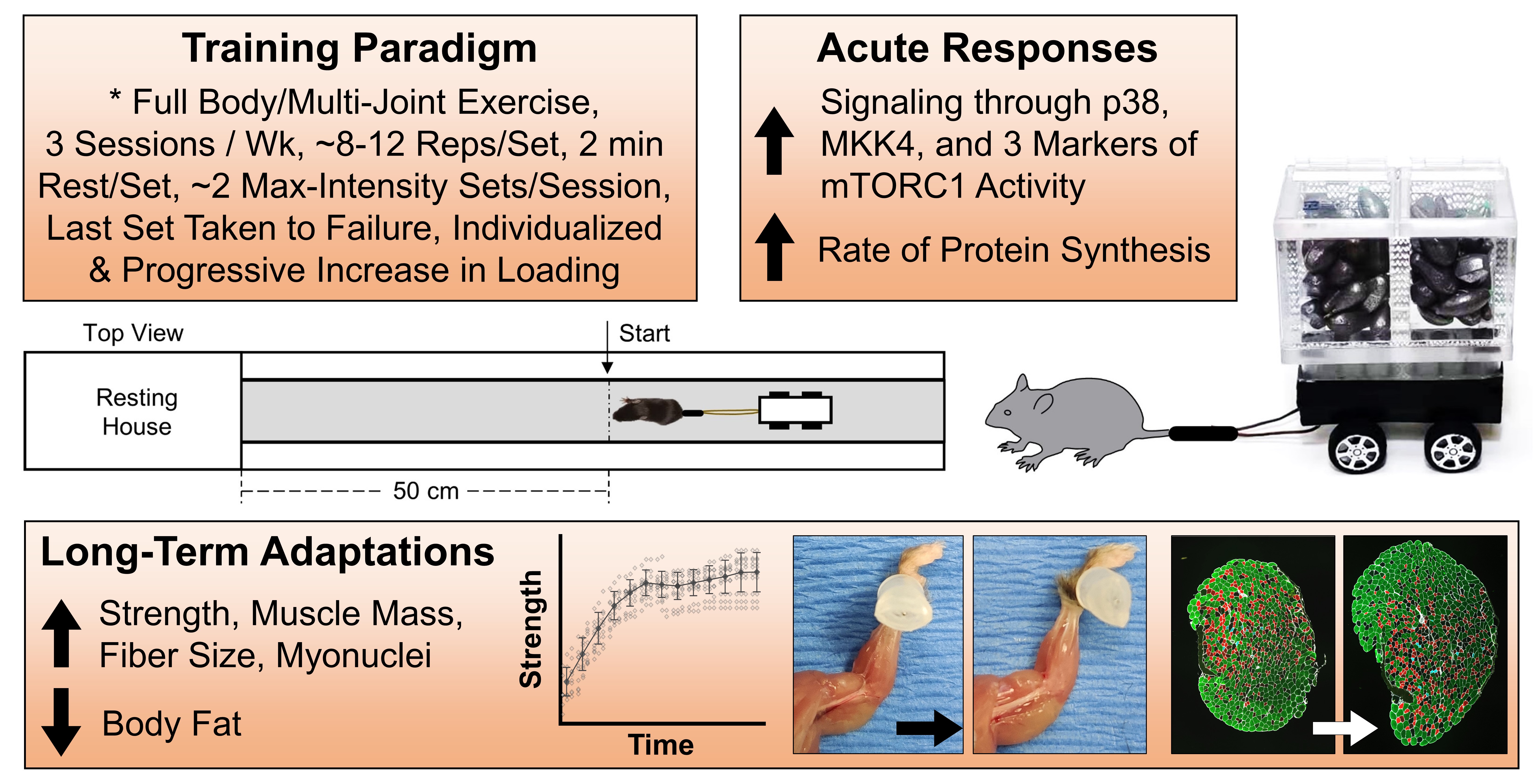

### Uncropped Western Blots

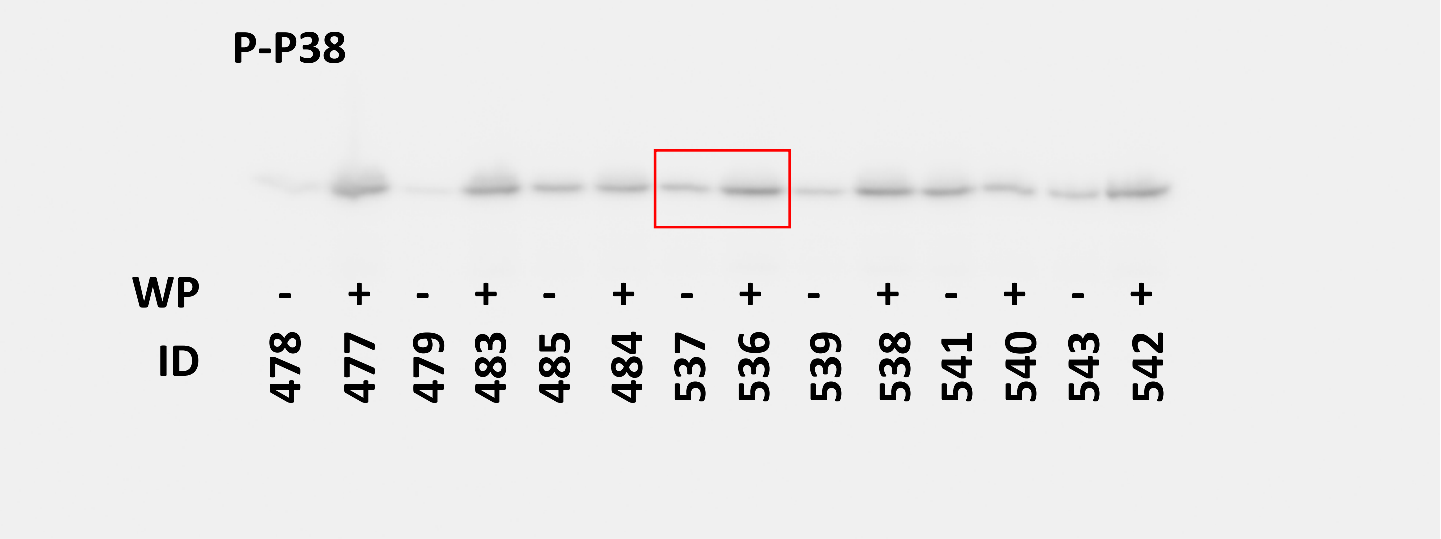

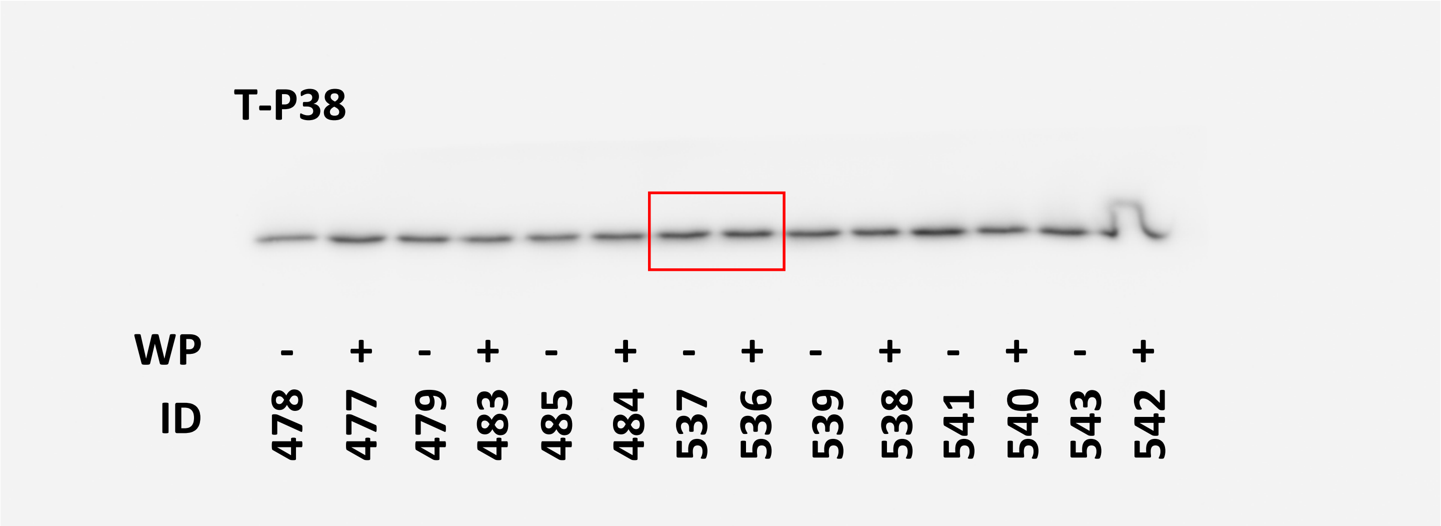

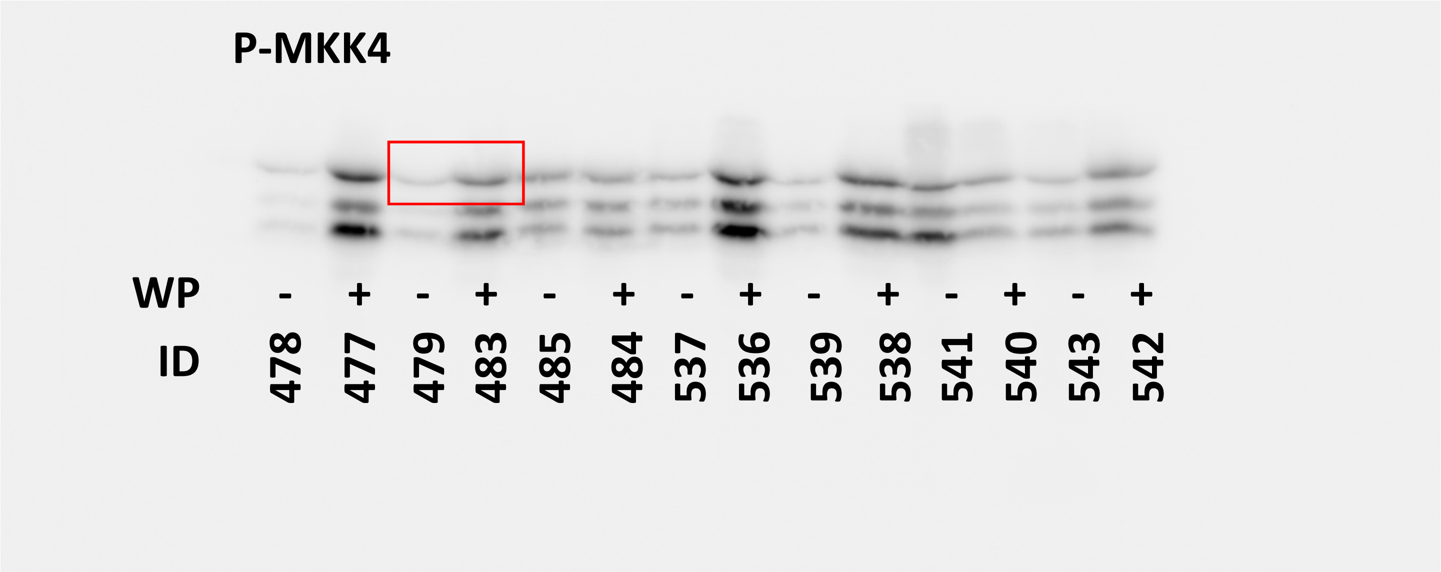

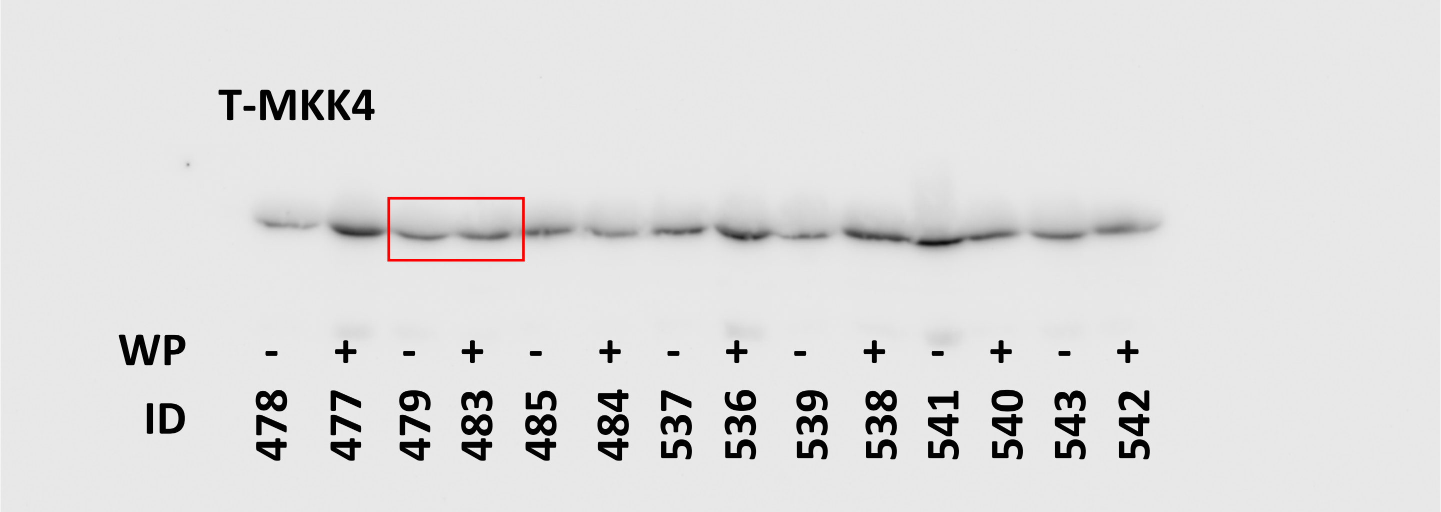

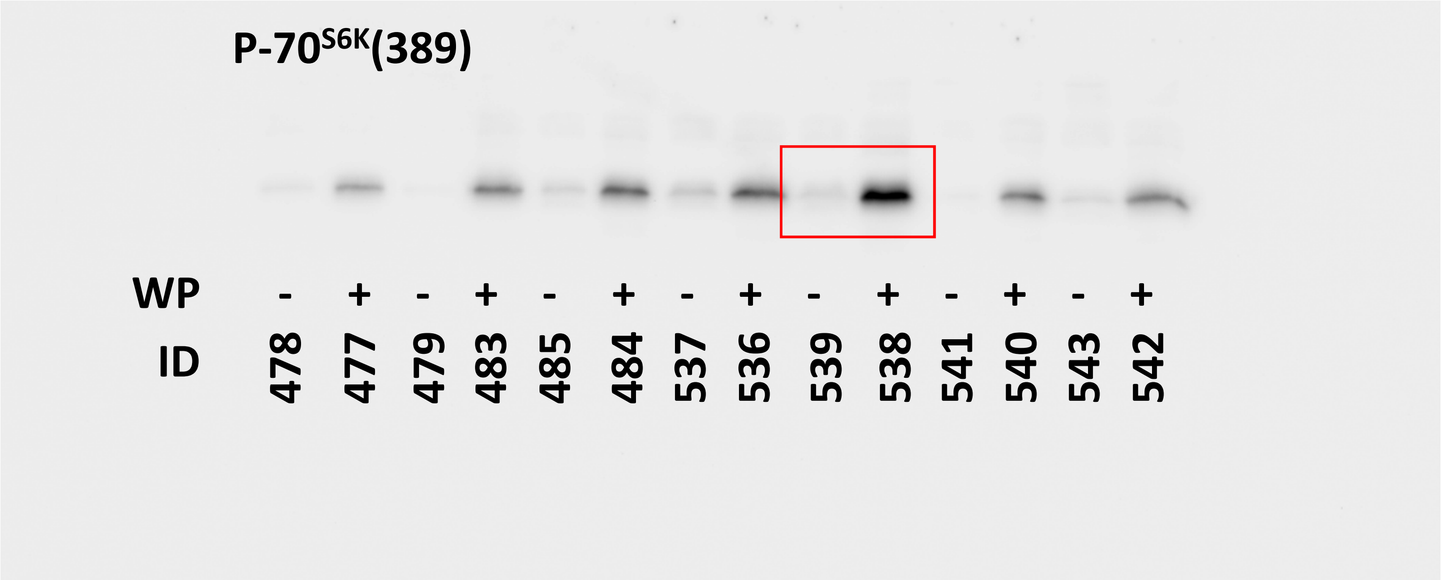

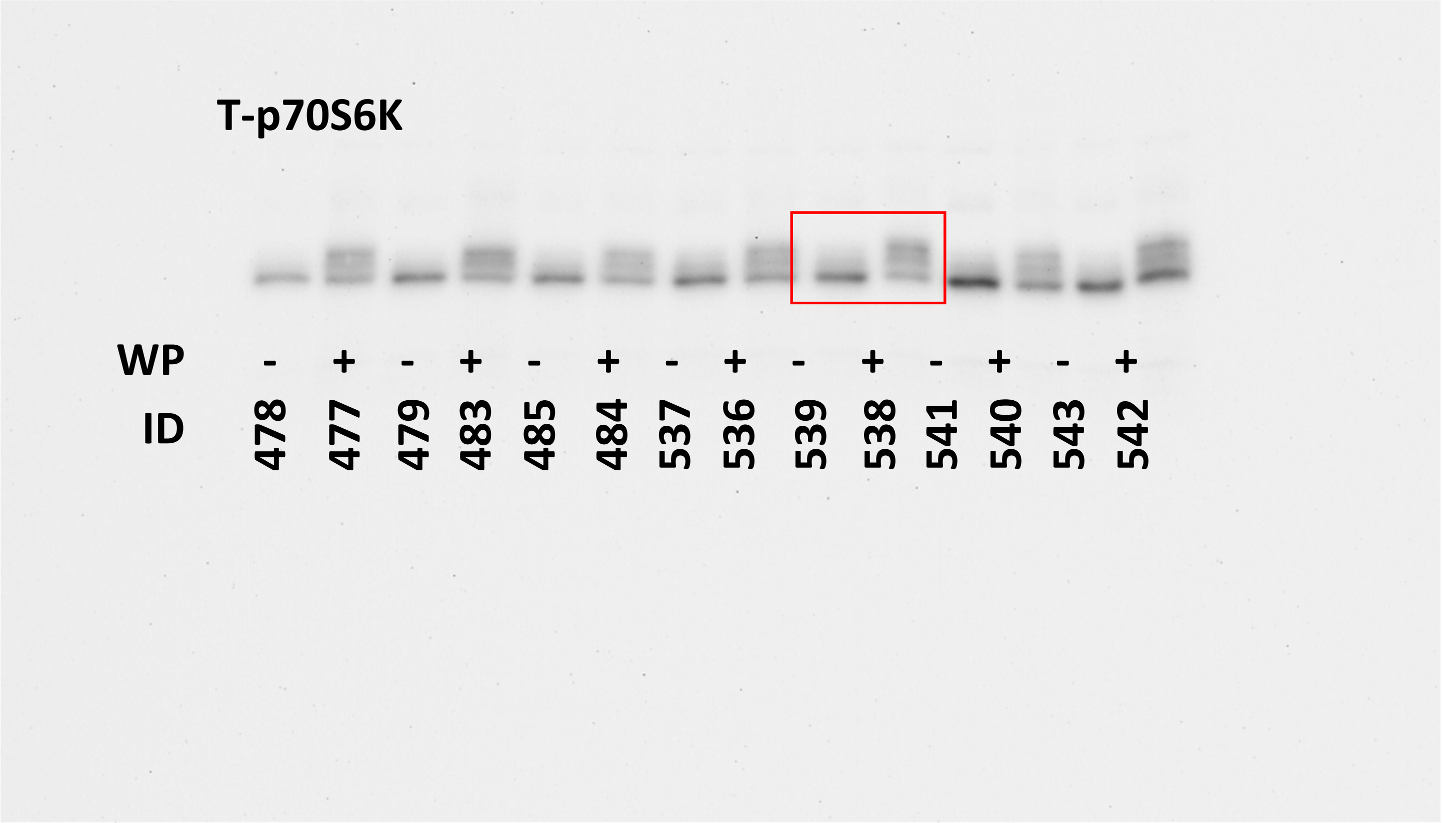

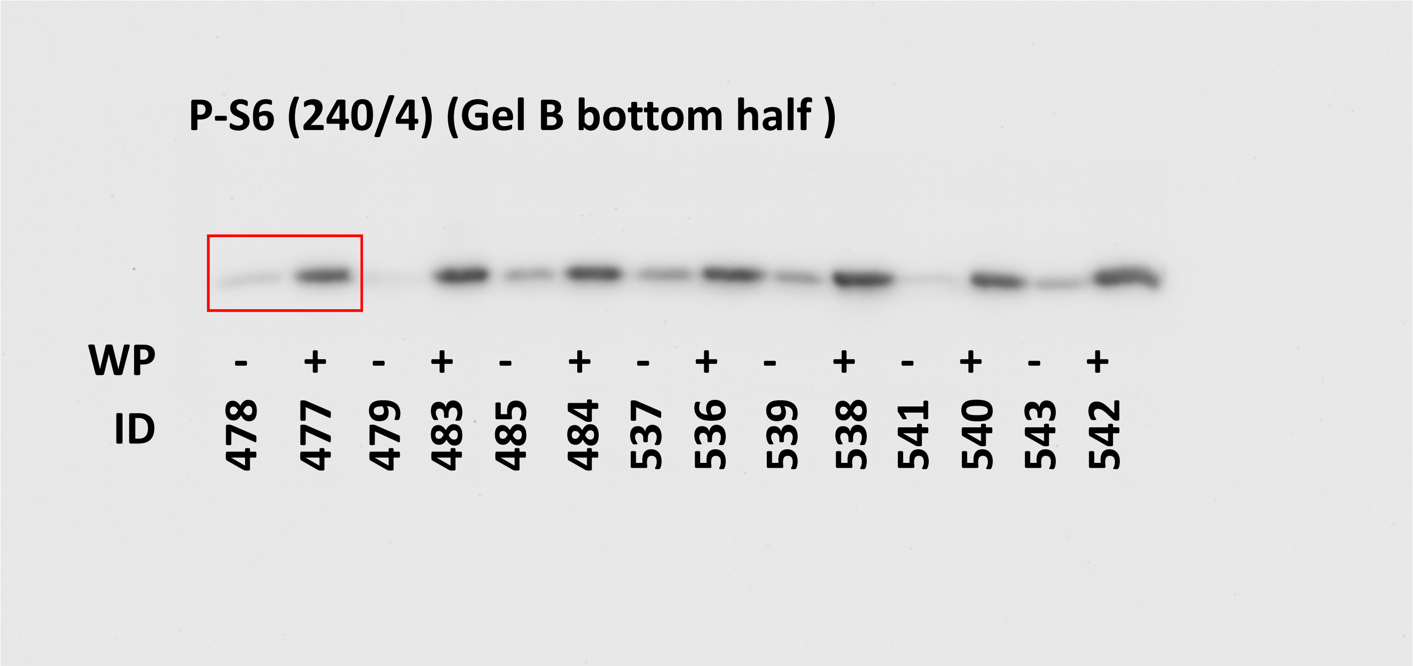

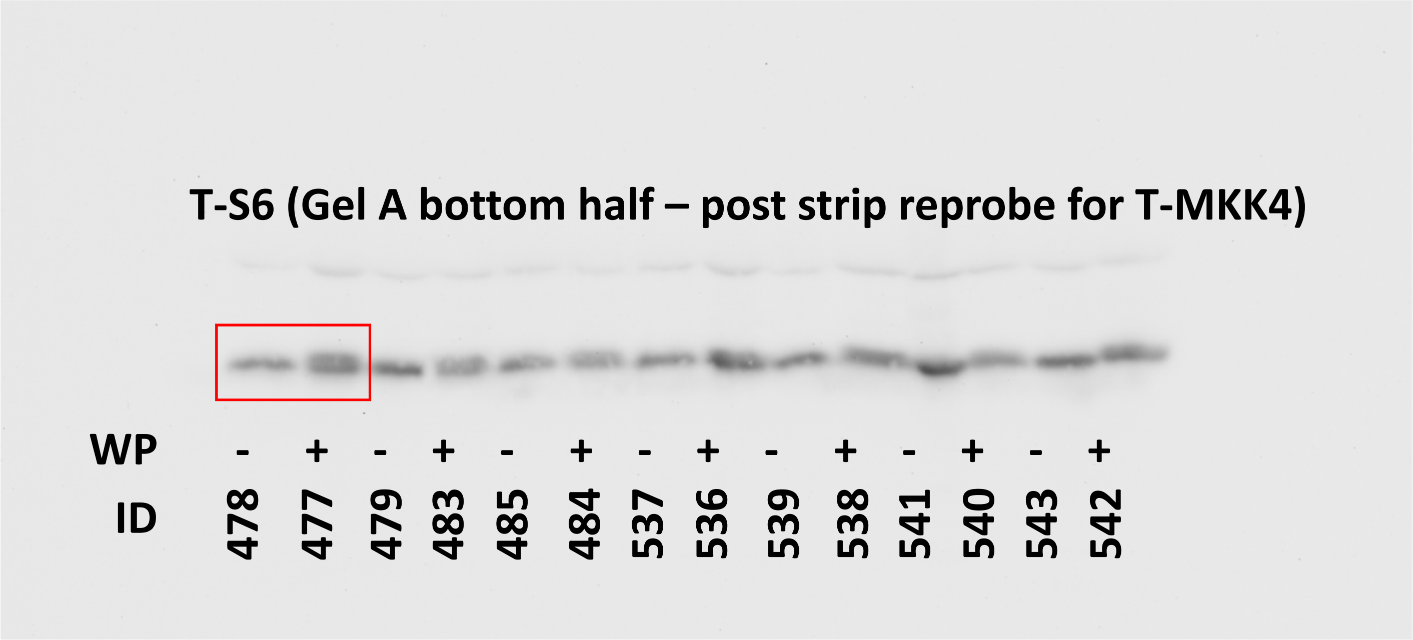

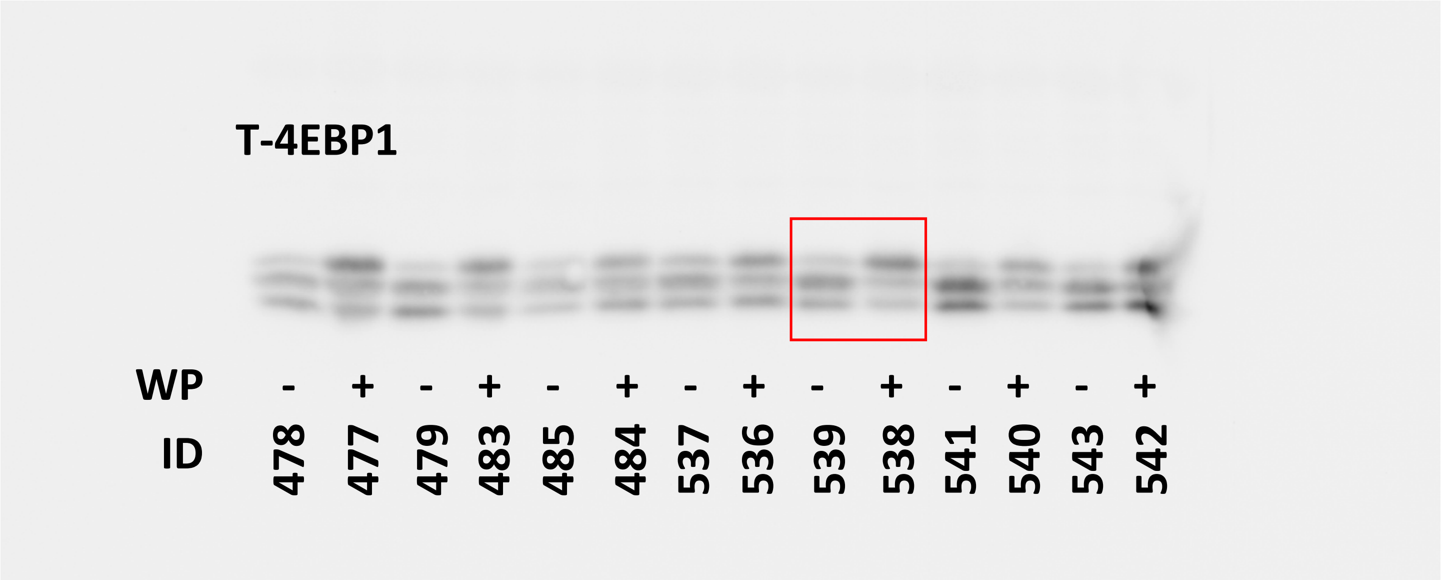

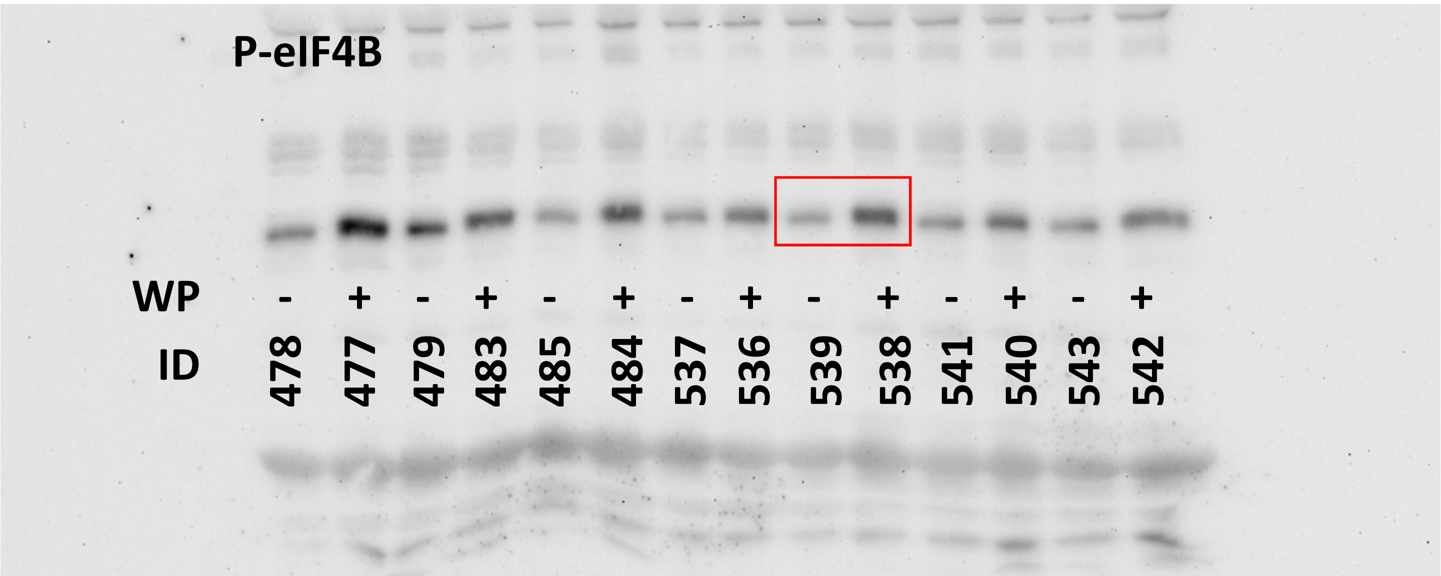

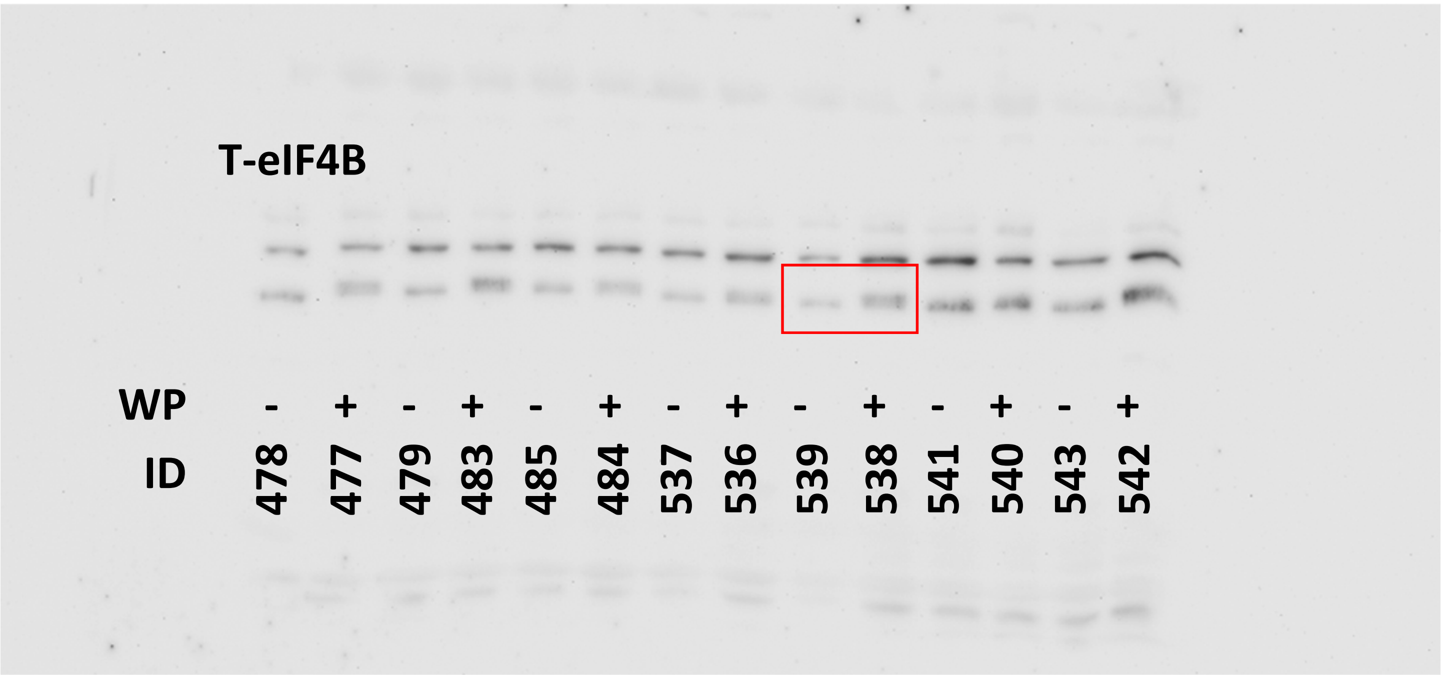

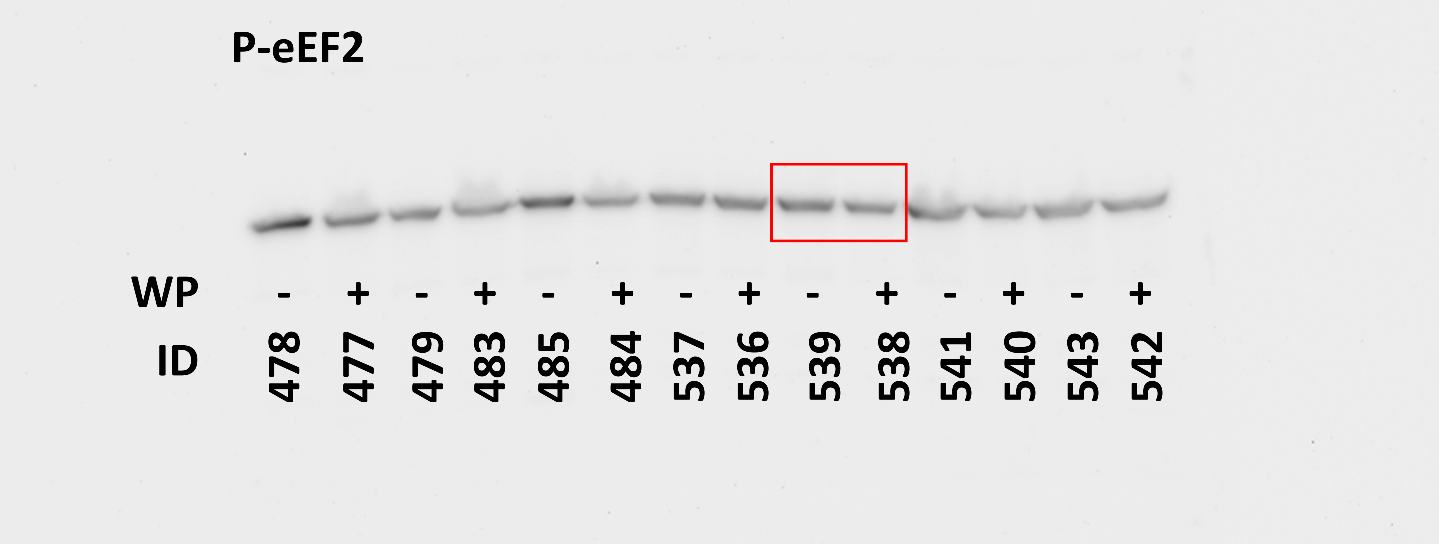

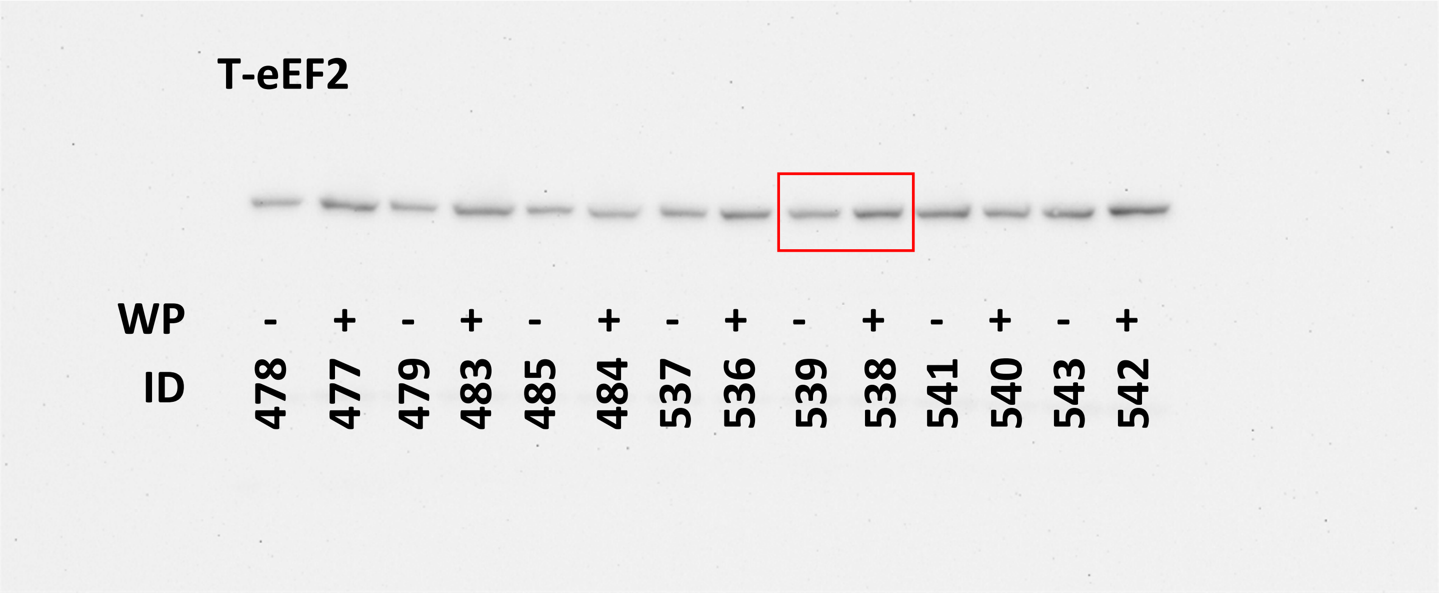

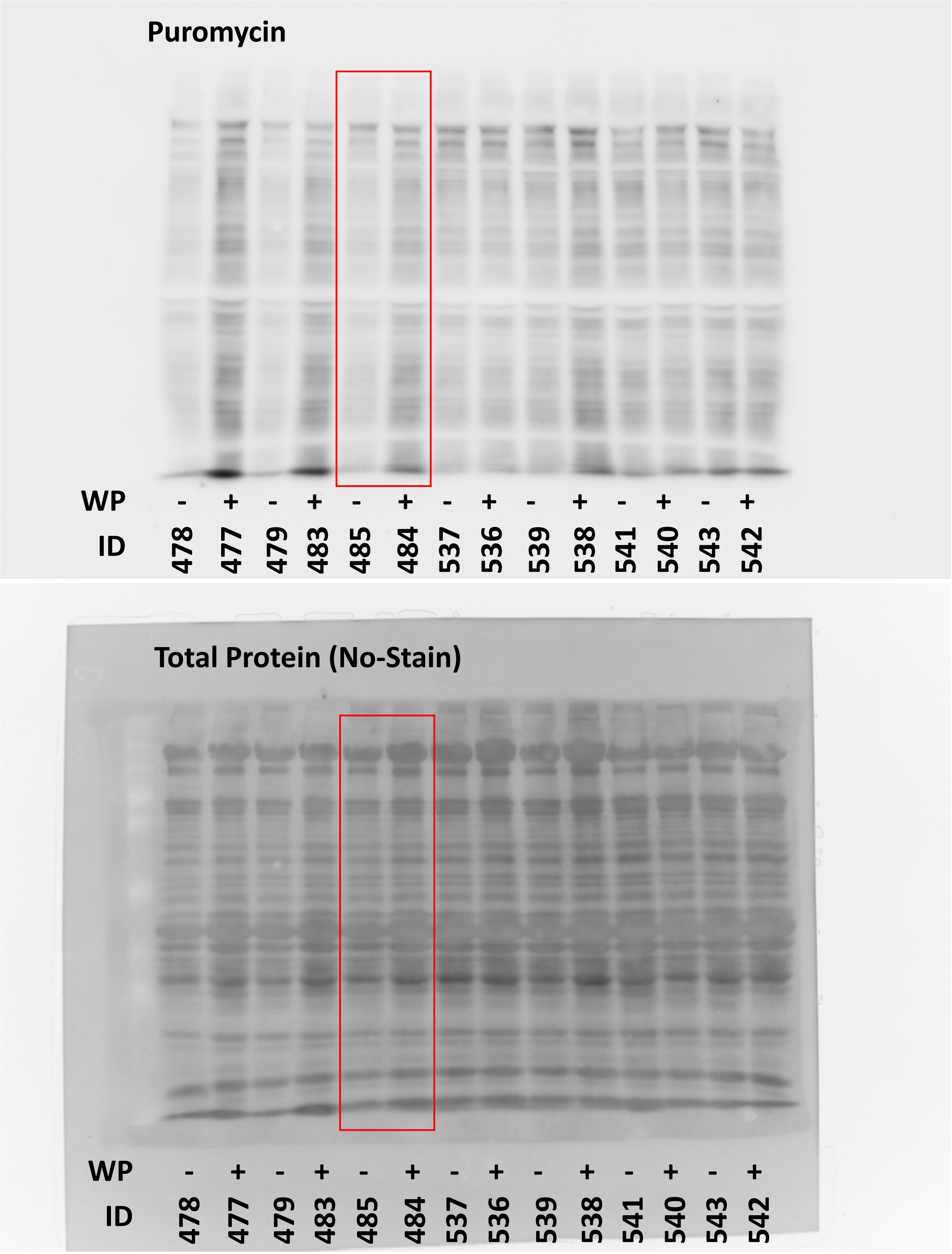

### Whole Cross-Section Images Merged.tif

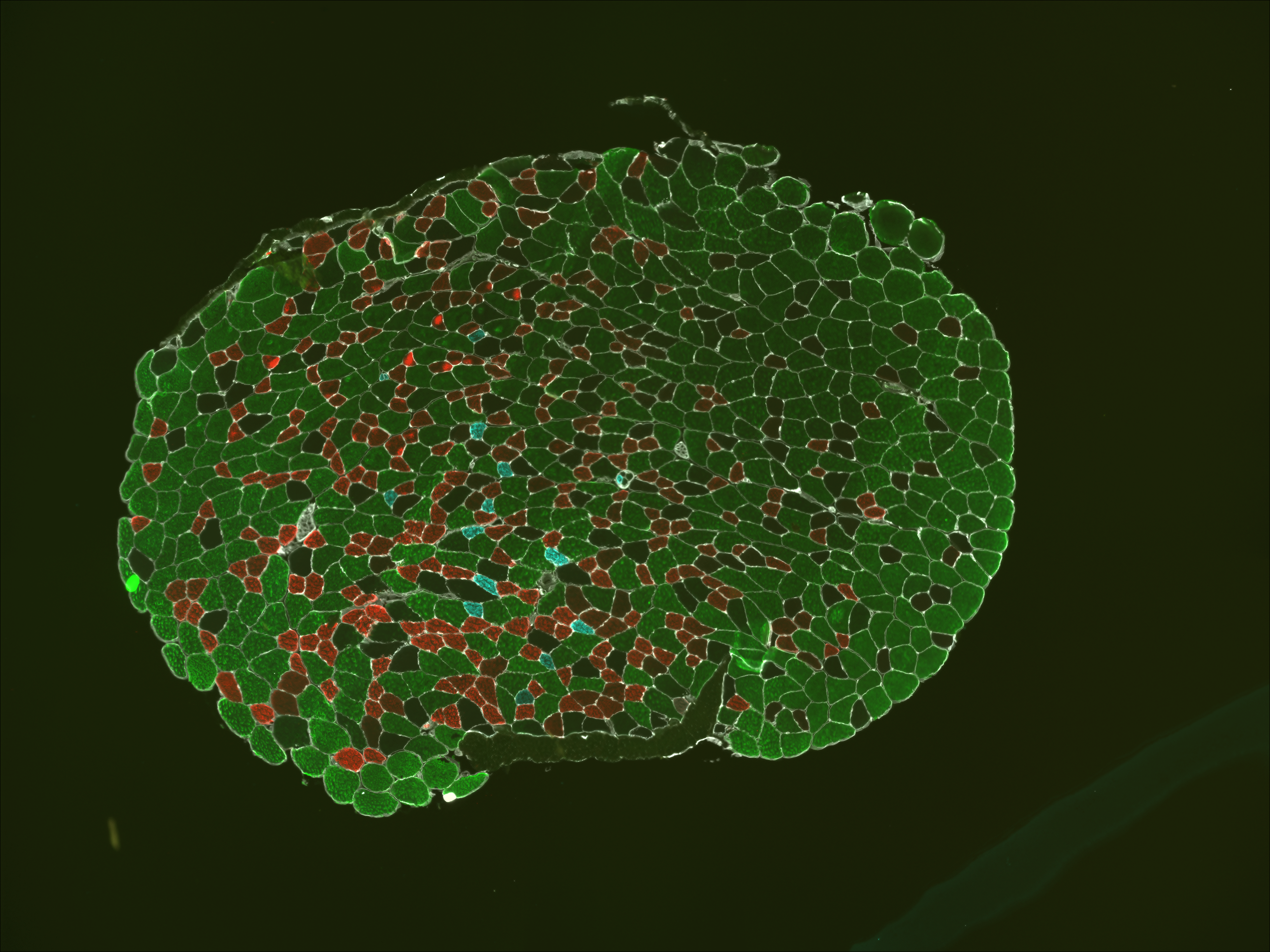

### Whole Cross-Section Images Merged.tif

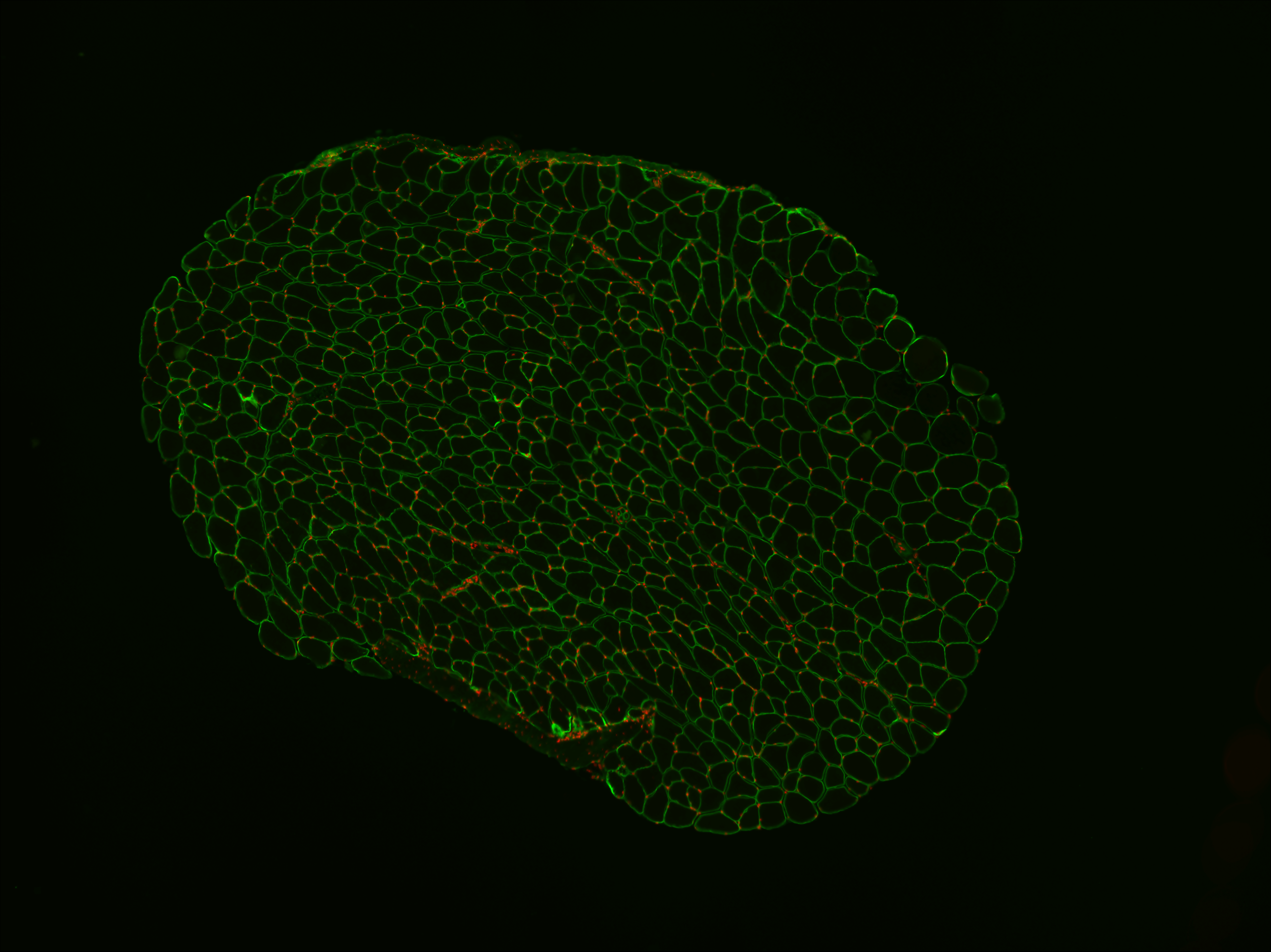
